# MAYOCTransformer: Masked-Attention for Yielding Comprehensive Semantic Segmentation of Retinal Optical Coherence Tomography Images using Transformer-based Neural Networks

**DOI:** 10.1101/2025.07.08.663601

**Authors:** Run Zhou Ye, Jenna Krivit, Gregor Reiter, Raymond Iezzi

**Affiliations:** Department of Ophthalmology, Mayo Clinic, Rochester, Minnesota, USA

**Keywords:** Optical coherence tomography, retina imaging, image segmentation, semantic segmentation, software tool, vision transformer

## Abstract

**Purpose:** Optical coherence tomography (OCT) is a widely used imaging modality in ophthalmology. Accurate semantic segmentation of these images is critical for both clinical and research applications, yet existing convolutional neural network (CNN)-based methods face challenges in generalizability and robustness. This study introduces MAYOCTransformer, the first transformer-based deep learning model for comprehensive semantic segmentation of OCT images, and evaluates its performance against CNN-based models.

**Methods:** A large dataset of 3,500 OCT images was manually segmented using an iterative deep learning-assisted workflow. The MAYOCTransformer model, based on the Mask2Former architecture, was trained and compared against CNN-based segmentation models, including U-Net, U-Net++, FPN, and DeepLabV3+. Comprehensive segmentation tasks included 10 retinal layer segmentation, choroid stroma and vessel segmentation, and the identification of 9 types of discrete pathological findings including intraretinal fluid (IRF), subretinal fluid (SRF), pigment epithelial detachment (PED), subretinal hyper-reflective material (SHRM), intraretinal hyper-reflective foci, and reticular pseudodrusen. Model performance was evaluated using the Dice similarity coefficient (DSC) on a hold-out test set with five-fold cross-validation. Additional validation was performed using external datasets, open-source segmentation models, and a randomized blinded expert evaluation.

**Results:** MAYOCTransformer outperformed CNN-based models in most segmentation tasks. Choroid segmentation performance was comparable between MAYOCTransformer and CNN models. External validation demonstrated the model’s generalizability, achieving higher DSC scores than publicly available segmentation models. A blinded expert evaluation showed that MAYOCTransformer’s segmentation was non-inferior to manual annotations.

**Conclusion:** MAYOCTransformer provides improved segmentation performance over CNN-based models. Its ability to generalize to external datasets suggests potential applicability in clinical and research settings.

## INTRODUCTION

Optical coherence tomography (OCT) has become an essential imaging modality in ophthalmology. The ability to accurately segment retinal layers and pathological findings in OCT images is crucial for both clinical decision-making and research applications. Manual segmentation by expert graders remains the gold standard but is time-consuming and subject to inter-observer variability. Automated segmentation methods offer the potential to enhance efficiency, reproducibility, and scalability in analyzing large-scale OCT datasets. However, achieving robust and generalizable segmentation remains a challenge due to variations in image quality, acquisition protocols, and the diverse range of retinal pathologies.

Convolutional neural networks (CNNs) have been widely employed for semantic segmentation of medical images, including OCT scans. U-Net [1] and its variants have demonstrated strong performance in delineating retinal layers and detecting pathological features such as intraretinal fluid (IRF), subretinal fluid (SRF) [2–5], and pigment epithelial detachments (PED) [6, 7]. CNN-based models have also been used for choroidal segmentation [8–13]. Despite these advancements, CNNs are usually limited by their local receptive fields, which can impede their ability to capture long-range dependencies [14], particularly in complex structures such as the choroid and epiretinal membrane (ERM).

Recent developments in transformer-based architectures have introduced new possibilities for medical image segmentation. Transformers, originally designed for natural language processing, have been adapted for vision tasks through self-attention mechanisms, enabling models to capture both local and global contextual information [15]. Studies comparing CNNs and transformers for OCT segmentation suggest that transformer-based models exhibit improved generalizability, particularly for images outside the training distribution [16, 17]. This advantage is particularly relevant in clinical applications, where segmentation models must handle images from different imaging systems and disease states without significant performance degradation.

In this study, we introduce **MAYOCTransformer**, **M**asked-**A**ttention for **Y**ielding Comprehensive Semantic Segmentation of Retinal **O**ptical **C**oherence **T**omography Images using **Transformer**-based Neural Networks. The model employs the masked-attention mask transformer (Mask2Former) architecture [18], which integrates multi-scale attention mechanisms to enhance feature representation across different anatomical structures. To benchmark MAYOCTransformer’s performance, we compare it against state-of-the-art CNN-based models, including U-Net [1], U-Net++ [19], FPN [20], and DeepLabV3+ [21], using a standardized training dataset curated from a large institutional image repository.

To further improve the efficiency of manual annotation, we developed OCTSegTool, a user-friendly software tool that facilitates iterative semi-automated segmentation. This tool enables an interactive deep learning-assisted workflow, where preliminary segmentation masks are refined through multiple iterations of human correction and model inference.

In addition to evaluating segmentation performance on a hold-out test set, we assess MAYOCTransformer’s generalizability using external OCT datasets with expert segmentation labels. Comparisons with publicly available segmentation models, such as ReLayNet [22] for retinal layers and Choroidalyser [8] for choroidal segmentation, provide insights into the model’s robustness across different imaging conditions [14,15]. Finally, we conduct a randomized blinded evaluation with surgical retina specialists, comparing MAYOCTransformer’s outputs against manual ground-truth annotations to assess clinical usability.

## METHODS

### Retina OCT image database

To develop robust semantic segmentation models that are generalizable for a wide variety of imaging conditions and pathologies, we first created an image database that contains all Heyex2 OCT slices stored in our ophthalmic image management system. Original OCT files stored in the DICOM (Digital Imaging and Communications in Medicine) format were converted to JPG images for ease of downstream model development. Cross-sectional OCT images were first separated from unrelated en-face images stored in the same image management system (*e.g.*, infrared images of the retina, fluorescein angiography, and indocyanine green (ICG) angiography images) and then subclassified into volumetric scans and radial scans using the original DICOM metadata tags. Subsequently, we used this image data pool to randomly sample a subset of 3,500 representative images for manual semantic segmentation and model training.

### Iterative method for semi-automated manual semantic segmentation

In order to accelerate the manual creation of semantic segmentation masks, we developed a pipeline and software tools to allow for iterative, deep-learning model assisted labelling (**Figure 1**). Our workflow began with randomly selecting a small subset of OCT images from the training set and performing manual labelling using our software tool (described in the following section). These image-mask pairs were then used to train a preliminary semantic segmentation model.

**Figure 1:**
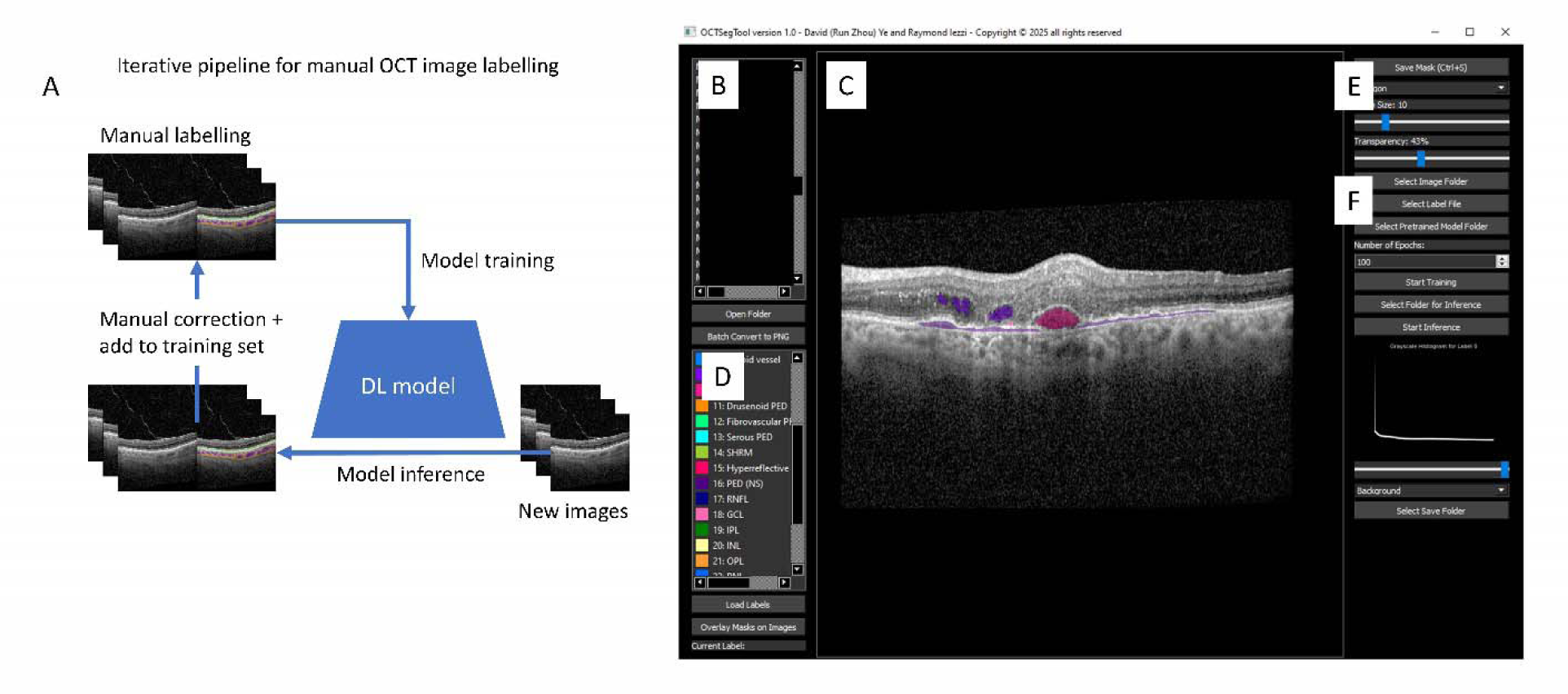
Overview of the iterative, deep learning–assisted labeling workflow and the OCTSegTool interface. Schematic of the iterative deep learning-assisted segmentation workflow used to generate manual labels for training(A). Main visualization panel of the OCTSegTool software (B). Segmentation mask overlay for the corresponding OCT image (C). Side panel listing all segmentation labels (D). Toolbar for brush size adjustment, polygon tool selection, transparency control (E). Integrated model training and inference interface, enabling users to train transformer-based models and perform segmentation directly from the software GUI (F).

This preliminary model was subsequently employed to generate new image-mask pairs with a new subset of OCT images taken from the training dataset, which would then undergo manual correction using our software tool and be added to our initial image-mask pairs. We iterated upon this pipeline until all images from the training dataset have been labelled.

### Development of the OCTSegTool

OCTSegTool is a user-friendly software tool that we created for iterative manual labeling and deep-learning model training specifically aimed at the semantic segmentation of fundus OCT images. To facilitate the use of our tool, we compiled our program into one single executable file that includes all dependency packages, without any installation needed. Developed in Python 3.8, OCTSegTool employs the PyQt5 library for its graphical user interface (GUI), facilitating loading of images, manual annotation, and saving of trained models and predicted segmentation masks. Image loading, augmentation, and preprocessing were implemented using Pillow, Numpy, OpenCV, and torchvision. For deep learning functionalities of model training and inference, the PyTorch library was utilized.

### OCT image segmentation classes

In this work, we aimed to create a set of deep-learning models that collectively generate accurate semantic segmentation masks for both normal retinal anatomical structures as well as discrete pathological findings. We trained a total of four models: an RPE model (for the segmentation of RPE and the outer segment of the photoreceptor layer); a retinal layer model (for the segmentation of all other retinal layers); a choroid model (for the segmentation of choroid vessels and stroma); and a discrete findings model (for the identification of abnormal findings in the retina). In addition, we also trained CNN-based models (including DeepLabV3+, FPN, U-Net, and U-Net++) using the same training dataset for comparison.

### OCT segmentation of retinal layers

In this study, we followed a similar retinal layer semantic segmentation definition as reported previously for retinal sublayer labelling [23–27] with a few adjustments. First, instead of identifying the boundary between retinal sublayers, we performed segmentation of the layers themselves as in [28]. This method of image labelling not only allows for a more direct measurement of, for example, RPE layer thickness in the setting of an RPE detachment from the Bruch’s membrane but is also more directly compatible with outputs from a transformer-based neural network. Furthermore, there are limited reports on the automated identification of epiretinal membrane (ERM) in the literature [29, 30]. It is also widely known that the presence of ERM introduces significant segmentation errors and can render automated RNFL and GCL layer segmentations unusable [31, 32]. For these reasons, we have included ERM as a separate class in order to make our layer segmentation model robust in the presence of ERM.

In summary, the following layers were labelled for the training of our retinal sublayer segmentation models (**Figure 2A-B**): ERM (epiretinal membrane); RNFL (retinal nerve fiber layer), GCL (ganglion cell layer); IPL (inner plexiform layer); INL (inner nuclear layer); OPL (outer plexiform layer); ONL (outer nuclear layer); PR-IS (photoreceptor-inner segment, corresponding to the region of photoreceptors between the external limiting membrane [ELM] and the middle of the ellipsoid zone [EZ] band [33]); PR-OS (photoreceptor-outer segment, defined as the region of photoreceptors between the middle of the EZ band and the inner border of the RPE); and the RPE (retinal pigment epithelium).

**Figure 2:**
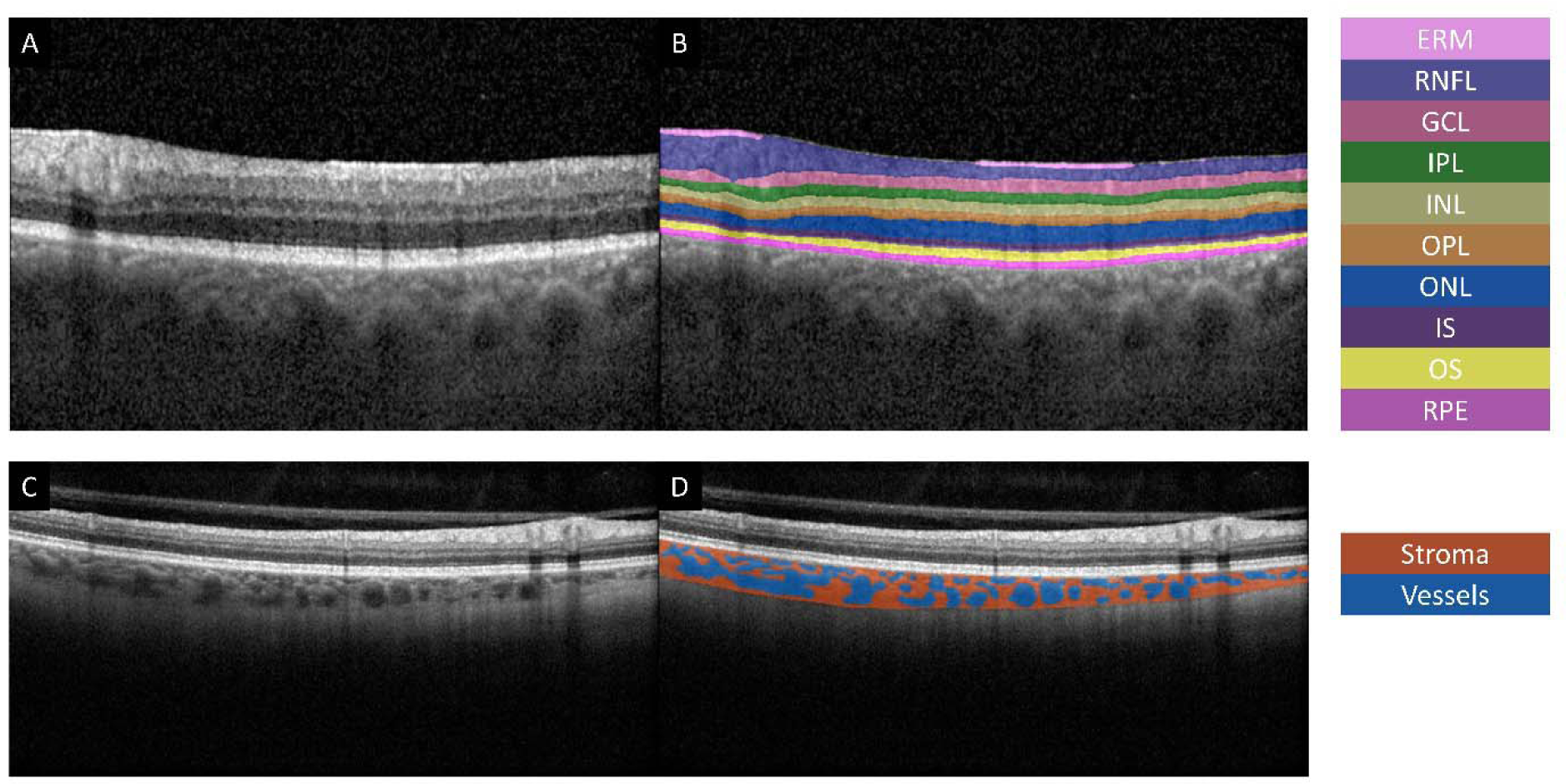
Macula OCT image (A) with corresponding semantic segmentation of retinal layers (B); labels include ERM, RNFL, GCL, IPL, INL, OPL, ONL, PR-IS, PR-OS, and RPE. Macula OCT image (C) with corresponding choroid stroma and vessel segmentation (D).

### OCT segmentation of choroid vessels and stroma

Both traditional machine learning algorithms [34–40] and deep-learning models have been developed for choroid segmentation. Notably, U-Net-based models [8–13] have been adapted for choroid vessel segmentation and fovea localization. In this work, we used the same definition for choroid vessel and stroma regions as previously reported: choroid is defined as the region between Bruch’s membrane and the choroid-scleral interface, and stroma is defined as choroid regions outside of vessel lumen (**Figure 2C-D**).

### OCT segmentation of intraretinal and subretinal fluid

Automated intraretinal fluid (IRF) and subretinal fluid (SRF) segmentation has been the focus of numerous previous reports. Existing methods for segmenting IRF and SRF include traditional machine-learning approaches [41–44] as well as deep-learning approaches [2–5]. Almost all existing deep-learning methodologies are based on CNNs. In the present study, we used the same definition for IRF and SRF as previously published (**Figure 3A-B**), with IRF defined as hyporeflective fluid regions within the neurosensory retina and SRF defined as fluid regions between the neurosensory retina and the RPE. In images with ERM, our IRF class excluded any hyporeflective regions between the ERM and the neurosensory retina (**Figure 3B and 3D**), which can easily be confused for IRF by CNN networks without explicit training. To our knowledge, this issue has not been addressed in previously published literature.

**Figure 3:**
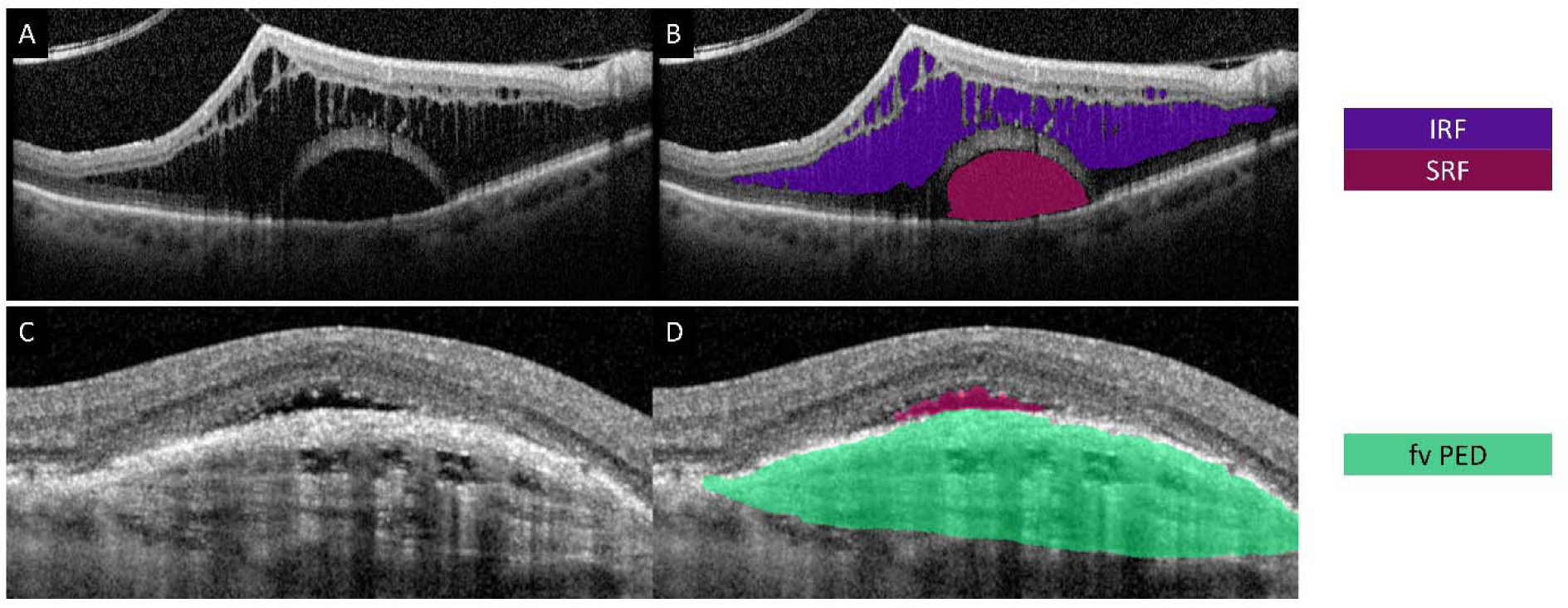
Macula OCT image (A) with corresponding semantic segmentation of IRF and SRF (B). Macula OCT image (C) with corresponding fibrovascular PED segmentation (D).

### OCT segmentation of pigment epithelium detachments

In this study, we subclassified pigment epithelium detachments (PEDs) into four main categories: fibrovascular PED (fv PED, **Figure 3C-D**), drusenoid PED (**Figure 4A-B**), serous PED (**Figure 4C-D**), and low-lying PED (**Figure 4E-F**). Automated segmentation of different types of PED have been previously described using manually designed algorithms [45–49] and deep-learning models [6, 7]. As previously reported, we defined fv PEDs as any irregular elevations of the RPE with fibrovascular tissue, which can take on the typical appearance of the “onion sign” [50].

**Figure 4:**
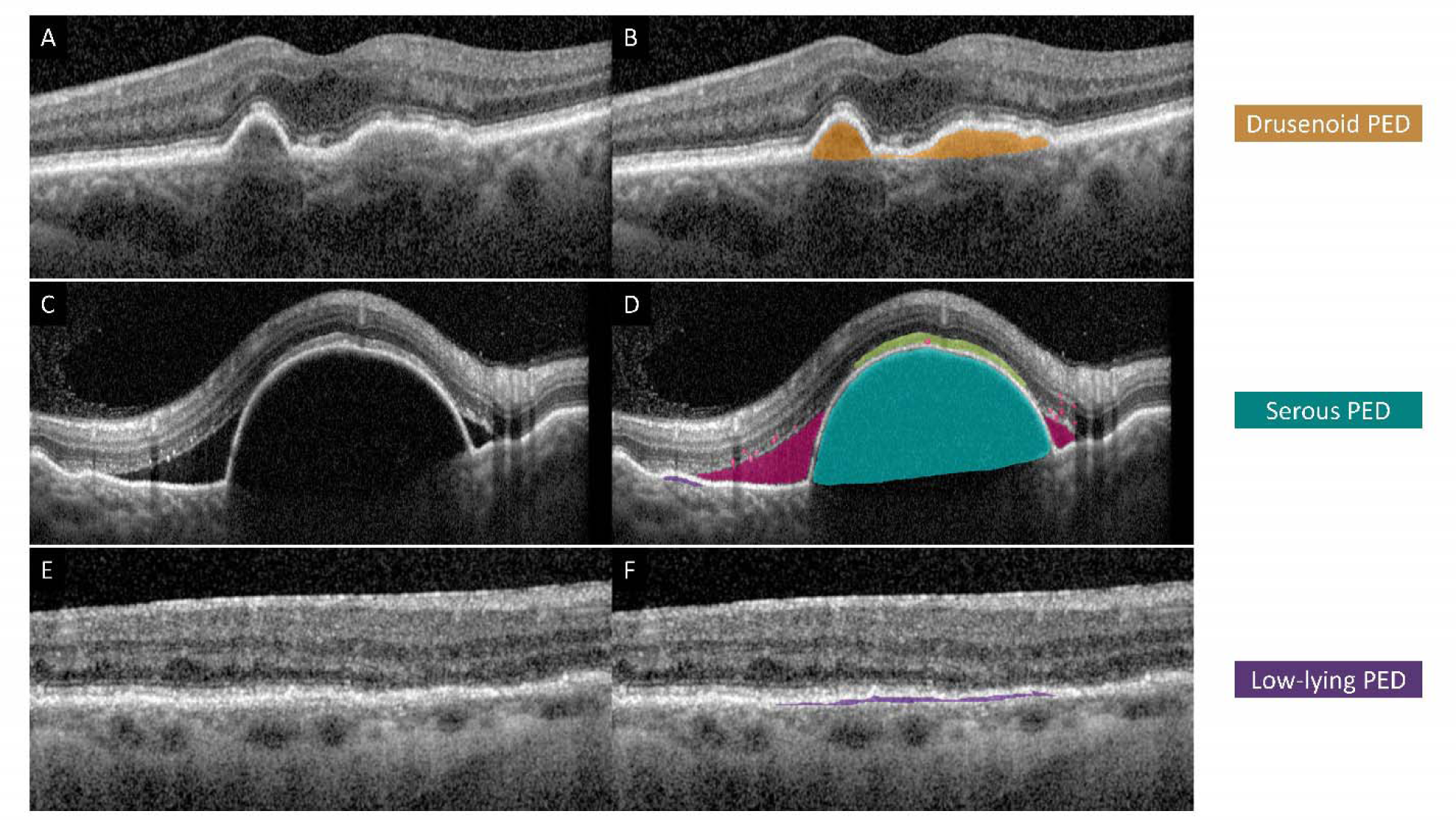
Subclassification of pigment epithelium detachments (PEDs). Macula OCT image (A, C, and E) with corresponding semantic segmentations of drusenoid PED (B), serous PED (D), and low-lying PED (F).

Drusenoid PEDs were defined as dome-shaped elevation of the RPE with relatively homogenously hyper-reflective core, as described in previous reports on automated segmentation [51–54]. In contrast, serous PEDs were defined as dome-shaped elevations of the RPE with homogenously hypo-reflective material. Additionally, low-lying PEDs were included as a separate category; these are defined as irregular and shallow elevations of the hyper-reflective RPE from the hyper-reflective Bruch’s membrane that creates the appearance of a “double layer” sign [55, 56].

### OCT segmentation of SHRM, reticular pseudodrusen, and hyper-reflective foci

Existing literature is scarce on automated segmentation of SHRM [7, 57, 58], reticular pseudodrusen [59], and hyper-reflective foci [60–62]. Consistent with previous reports, we defined SHRM as hyper-reflective regions between the neurosensory retina and the RPE (**Figure 5A-B**) and reticular pseudodrusen as small drusen-like deposits between the neurosensory retina as the RPE (**Figure 5C-D**). We defined hyper-reflective foci (HR foci) as any spot-shaped and highly hyper-reflective areas within the neurosensory retina (**Figure 5E-F**), including areas of RPE migration (**Figure 5G-H**).

**Figure 5:**
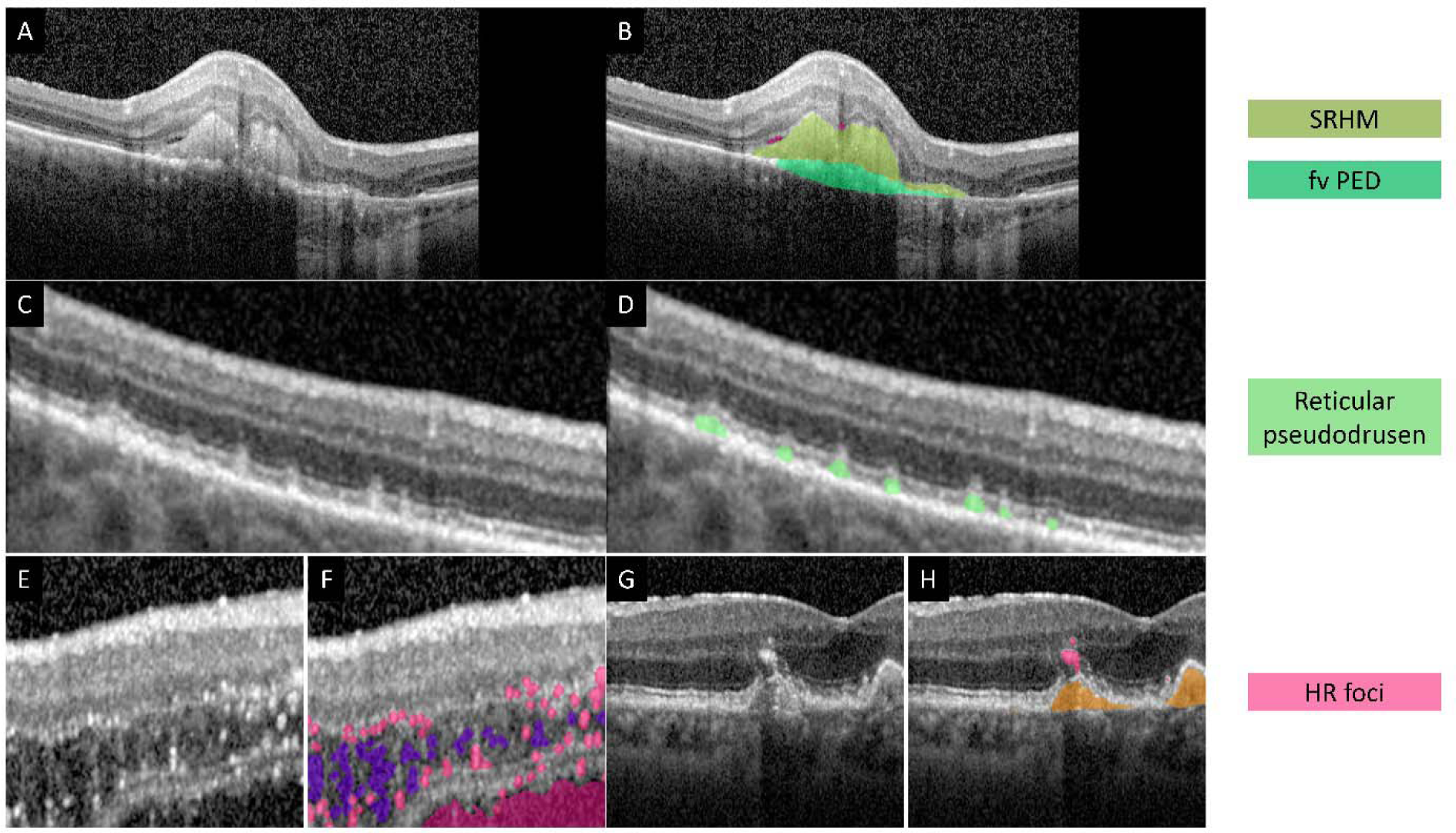
Segmentation of SHRM, reticular pseudodrusen, and hyper-reflective foci. OCT image (A) showing subretinal hyper-reflective material (SHRM) with corresponding segmentation (B). OCT image (C) showing reticular pseudodrusen between the retina and RPE with corresponding segmentation (D). Hyper-reflective foci (HR foci) visible as small bright spots within the retina € with corresponding segmentation (F). Example of RPE migration with HR foci pattern (G) with corresponding segmentation (H).

### Segmentation model training

Convolutional neural networks (CNNs), particularly the U-Net architecture [1], have been the standard for semantic segmentation in medical imaging. Nonetheless, with the introduction of vision transformers [15], recent advances have seen transformer-based networks surpass all types of CNNs in various image segmentation tasks, in part thanks to their ability to better capture global semantic dependencies as well as their better generalizability. For example, in the context of fundus photo classification [16] and OCT segmentation [17], CNN-based models showed acceptable performance for images in the training datasets; however, their accuracy drops significantly when given images that are out of the training distribution, while transformer-based models performed significantly better on images outside of the training dataset. Indeed, transformer models now dominate the state-of-the-art in image segmentation across different domains, as reviewed in [63] and [64].

In the present study, we employed the masked-attention mask transformer (Mask2Former) architecture [18] as our semantic segmentation model. We compared the Mask2Former architecture against the CNN architectures for OCT image segmentation, including the U-Net [1], Unet++ [19], FPN [20], and the DeepLabV3+ [21].

After creating the training image-mask pairs using the iterative labelling method, weights of all models were re-initialized and subsequently trained using a 0.8 : 0.1 : 0.1 training : validation : testing split with a 5-fold cross-validation scheme. Models were trained using the AdamW optimizer and binary cross entropy loss for 100 epochs with a learning rate of 0.00006 and a batch size of 1.

To improve the robustness and generalizability of the deep learning models, several data augmentation techniques were applied. These included random horizontal flipping of images with a probability of 0.5, scaling images within a range of 0.7 to 1.3, translating images within a range of 0.3 of the image size. All augmentations were implemented using the Albumentations library [65].

### Model validation against manual labelling variability

To measure manual labelling variability and compare manual variability against model segmentation error, we manually re-labelled 40, 20, and 20 images for discrete findings, retinal layers and choroid segmentation using images randomly selected from the hold-out test set. For manual re-labelling, we labelled regions of interest entirely by hand using the paint tool from our OCTSegTool software without any model assistance. The mean absolute pixel area difference of each region of interest is calculated between two versions of manual segmentation maps and between the model *vs.* manual segmentation maps.

### Model validation using external datasets

To further validate the generalizability of our models, we tested the accuracy of our model segmentation on public OCT datasets with expert segmentation labels. For retinal fluid segmentation, we tested model accuracy using the dataset by Ahmed et al. [66], which contains 1200 OCT images with corresponding groundtruth segmentation maps. For retinal layer segmentation, we tested model accuracy against the OCT5K dataset [67], which contains nearly 5000 macula OCT images with corresponding groundtruth segmentation of retinal sublaminae. For choroid stroma and vessel segmentation, there is no publicly available dataset to our knowledge; we therefore tested model performance using the OCT2017 dataset by Kermany et al. [68]; we further curated 17 publicly available swept-source OCT images and performed manual segmentation for choroid stroma as well as Sattler and Haller layer vessels without model assistance to test model generalizability to different types of OCT images.

### Comparison with open-source public models

To further compare our models against publicly available state-of-the art open-weight neural network models, we performed head-to-head comparisons of our models against the ReLayNet [22] and the Choroidalyzer [8] models (using the weights as provided by the authors) for layer and choroid segmentation using the OCT5K dataset and our curated swept-source OCT images, respectively, without any finetuning.

### Randomized blinded testing for segmentation model validation

To further assess the validity of model segmentation, we first randomly selected and manually labelled 83 new macula OCT images from new patients from our image pool. Model segmentation was then performed for the same images. We devised an online quiz-based randomized blinded testing using the Qualtrics platform whereby manual and model segmentation maps were shown in random order to a group of 2 retina specialists with 29 and 8 years of clinical experience, who were instructed to choose which segmentation map they prefer, without any prior knowledge of which image corresponds to the manual or model segmentation. For images containing multiple types of findings, we created a separate multiple-choice question for each finding, for a total of 349 questions. For each question (“Which segmentation is better?”), 3 multiple choices were presented: image “A”, image “B”, and “Same” (**Figure S1**).

For each question, segmentations were scored as follows: +0 point if the specialist chose the alternative segmentation, +1 point was given if the specialist chose “Same” as the answer, +2 if the specialist chose the same segmentation. Points from both specialists were added for each question and compared between model *vs.* manual segmentations. We further assessed inter-rater agreement by defining disagreement as specialists choosing opposite segmentations.

### Statistical analyses

All statistical analyses were performed using SPSS 20 and Python (version 3.8) with SciPy and NumPy libraries. Statistical significance was defined as p < 0.05. To compare segmentation performance between MAYOCTransformer and CNN-based models (DeepLabV3+, FPN, U-Net, U-Net++), we computed the Dice similarity coefficient (DSC) for each segmentation class.

Pairwise comparisons were conducted using Tukey’s pair-wise multiple comparisons test. For assessing the variability between manual labelling and model segmentation, we calculated the mean absolute area difference for each segmentation class. A two-sample t-test was used to compare variability in manual annotations against segmentation errors from MAYOCTransformer. To evaluate external dataset performance, the Dice coefficient was computed between ground-truth segmentation maps and model predictions.

## RESULTS

### Retina OCT image database

Using our ophthalmology image management system, we created a large cross-sectional OCT image database of over 18M images. These images included 16M volumetric and 2M radial cross-sectional images, which have been taken at Mayo Clinic, Rochester between July 3, 2008 and October 24, 2024, from a total of 41,820 patients. **Table S1** summarizes the baseline characteristics of our complete cross-sectional OCT image database. In brief, image widths ranged from 384 to 1152 pixels, with signal-to-noise ratio ranging from 0.36 to 2.32. There were no significant deviations in these image characteristics between the training subset and the entire image pool.

### OCTSegTool software

The OCTSegTool software offers a comprehensive graphical user interface (**Figure 1B-F**) designed to facilitate image segmentation tasks. The interface is divided into several sections: the main window for visualizing input images and corresponding masks (**Figure 1B and C**), a side panel for selecting and managing labels (**Figure 1D**), and a set of controls for adjusting brush size, transparency, and tool selection (**Figure 1E**). Additionally, the interface includes options for loading images, saving masks, and initiating batch conversions between image formats. A unique feature of our tool is its integration with deep learning model training and inferencing directly from the user interface, allowing users to select image folders, label files, and pretrained models (**Figure 1F**).

The general workflow when using OCTSegTool begins with loading the images and associated labels for segmentation. Users can manually annotate the images using the brush or polygon tools, with the option to toggle between different labels or adjust transparency to better view underlying image details. The tool supports automatic mask generation through deep learning, utilizing transformer-based semantic segmentation models. Users can initiate model training directly from the interface; the trained model can then be used for inference on new images, with the resulting segmentation masks saved automatically.

Throughout the training process, model weights can be saved at the end of each epoch if they show improved performance, ensuring that users can easily load and continue training from the best-performing model. This feature also supports transfer learning, where a pretrained model can be fine-tuned on a different dataset or for a different segmentation task.

### Automated segmentation of retinal layers, choroid vessels and stroma

As shown in **Table 1** and **Figure S2**, the MAYOCTransformer model outperformed other CNN-based models in semantic segmentation of most of retinal layers. This improvement was statistically significant for the RNFL, GCL, and ERM classes, but did not reach statistical significance for other layers. For PR-OS and RPE segmentation, our U-Net model performed marginally better than the MAYOCTransformer model.

**Table 1:** Performance of segmentation models on hold-out test dataset with 5-fold cross-validation

| Segmentation class | MAYOCTransformer | DeepLabV3+ | FPN | U-Net | U-Net++ |
| --- | --- | --- | --- | --- | --- |
| **Layers** |  |  |  |  |  |
| **RNFL** | **0.93 ± 0.01** | 0.85 ± 0.05* | 0.88 ± 0.01* | 0.87 ± 0.04* | 0.86 ± 0.02* |
| **GCL** | **0.92 ± 0.01** | 0.83 ± 0.04* | 0.86 ± 0.02* | 0.86 ± 0.02* | 0.86 ± 0.02* |
| **IPL** | **0.89 ± 0.01** | 0.83 ± 0.04* | 0.87 ± 0.02 | 0.87 ± 0.01 | 0.86 ± 0.02 |
| **INL** | **0.88 ± 0.02** | 0.84 ± 0.04 | 0.87 ± 0.03 | 0.87 ± 0.03 | 0.86 ± 0.03 |
| **OPL** | **0.88 ± 0.04** | 0.85 ± 0.03 | 0.86 ± 0.03 | 0.87 ± 0.02 | 0.86 ± 0.03 |
| **ONL** | **0.93 ± 0.01** | 0.89 ± 0.03 | 0.89 ± 0.04 | 0.90 ± 0.03 | 0.89 ± 0.03 |
| **PR IS** | **0.84 ± 0.01** | 0.82 ± 0.03 | 0.82 ± 0.05 | 0.83 ± 0.04 | 0.83 ± 0.03 |
| **PR OS** | 0.88 ± 0.008 | 0.87 ± 0.03 | 0.87 ± 0.02 | **0.89 ± 0.01** | 0.88 ± 0.02 |
| **RPE** | 0.87 ± 0.01 | 0.86 ± 0.02 | 0.86 ± 0.02 | **0.87 ± 0.04** | 0.87 ± 0.02 |
| **ERM** | **0.81 ± 0.02** | 0.50 ± 0.1** | 0.47 ± 0.1** | 0.48 ± 0.07** | 0.49 ± 0.08** |
| **Choroid** |  |  |  |  |  |
| **Stroma** | **0.91 ± 0.001** | 0.87 ± 0.003** | 0.88 ± 0.003** | 0.89 ± 0.005** | 0.90 ± 0.002 |
| **Vessel** | **0.92 ± 0.001** | 0.89 ± 0.005* | 0.90 ± 0.007 | 0.90 ± 0.009* | 0.92 ± 0.005 |
| **Discrete findings** |  |  |  |  |  |
| **IRF** | **0.84 ± 0.003** | 0.80 ± 0.001 | 0.74 ± 0.02 | 0.77 ± 0.002 | 0.77 ± 0.002 |
| **SRF** | 0.84 ± 0.008 | 0.81 ± 0.003 | **0.87 ± 0.001** | 0.79 ± 0.003 | 0.79 ± 0.02 |
| **Drusenoid PED** | **0.79 ± 0.02** | 0.69 ± 0.02 | 0.64 ± 0.02 | 0.71 ± 0.02 | 0.58 ± 0.03 |
| **Fv PED** | **0.64 ± 0.03** | 0.48 ± 0.02* | 0.36 ± 0.002** | 0.34 ± 0.008** | 0.31 ± 0.005** |
| **Serous PED** | **0.69 ± 0.001** | 0.58 ± 0.003 | 0.66 ± 0.05 | 0.62 ± 0.01 | 0.60 ± 0.001 |
| **Low-lying PED** | **0.66 ± 0.03** | 0.59 ± 0.001 | 0.54 ± 0.003 | 0.59 ± 0.001 | 0.60 ± 0.001 |
| **SHRM** | **0.78 ± 0.009** | 0.47 ± 0.06 | 0.58 ± 0.02 | 0.21 ± 0.03** | 0.37 ± 0.03** |
| **HR foci** | **0.64 ± 0.03** | 0.51 ± 0.001 | 0.51 ± 0.001 | 0.52 ± 0.001 | 0.57 ± 0.001 |
| **Reticular pseudodrusen** | **0.58 ± 0.02** | 0.43 ± 0.001 | 0.38 ± 0.003* | 0.35 ± 0.009** | 0.40 ± 0.001* |

Values are mean ± variance. \* p-value < 0.05, \*\* p-value < 0.01. p-values are for Tukey's pairwise multiple comparisons against results for MAYOCTransformer. Bolded values represent the highest averages for a given class.

For choroid stroma and vessel segmentation, the MAYOCTransformer model performed marginally better compared to all other CNN-based models (**Table 1** and **Figure S3**). **Figure 6** shows examples of layer segmentation and choroid segmentation of testing OCT images.

**Figure 6:**
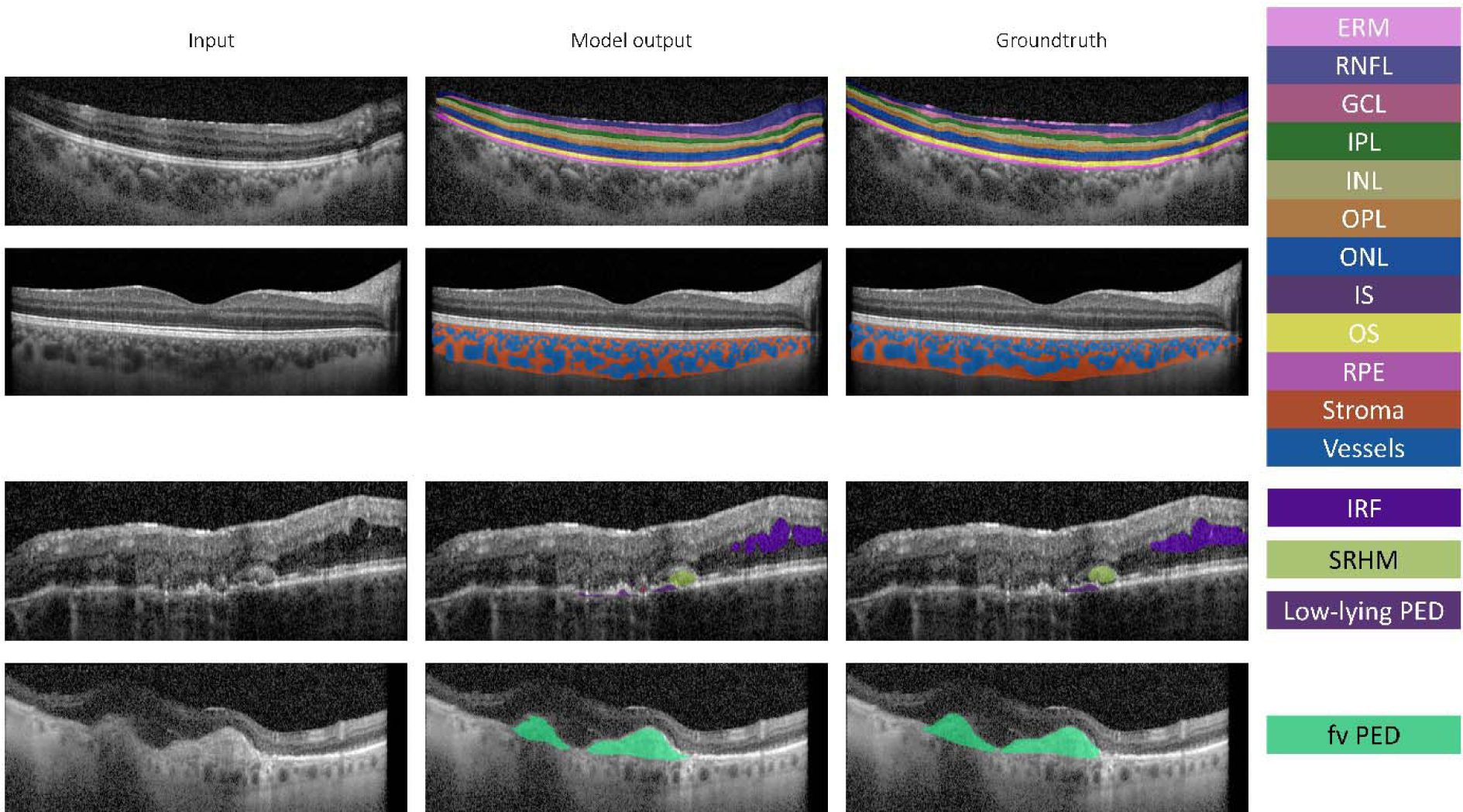
Comparison of MAYOCTransformer segmentation output with groundtruth segmentation masks of retinal layers, IRF, SHRM, low-lying PED, and fv PED. Input OCT images (first column) with model segmentation (second column), and groundtruth segmentation (third column).

### Automated segmentation of IRF, SRF, PED, SHRM, reticular pseudodrusen, and hyper-reflective foci

As shown in **Table 1** and **Figures 6-7**, our models achieved good semantic segmentation accuracy for discrete OCT findings in a hold-out testing dataset. In general, the MAYOCTransformer model outperformed all tested CNN-based model architectures that have been trained using the same training dataset, including DeepLabV3+, FPN, U-Net, and U-Net++ (**Figure S4**). However, these differences in performance were statistically significant only for fv PED, SHRM, and reticular pseudodrusen and did not reach statistical significance for other types of discrete findings. Hence, further model assessments were performed only with the MAYOCTransformer.

**Figure 7:**
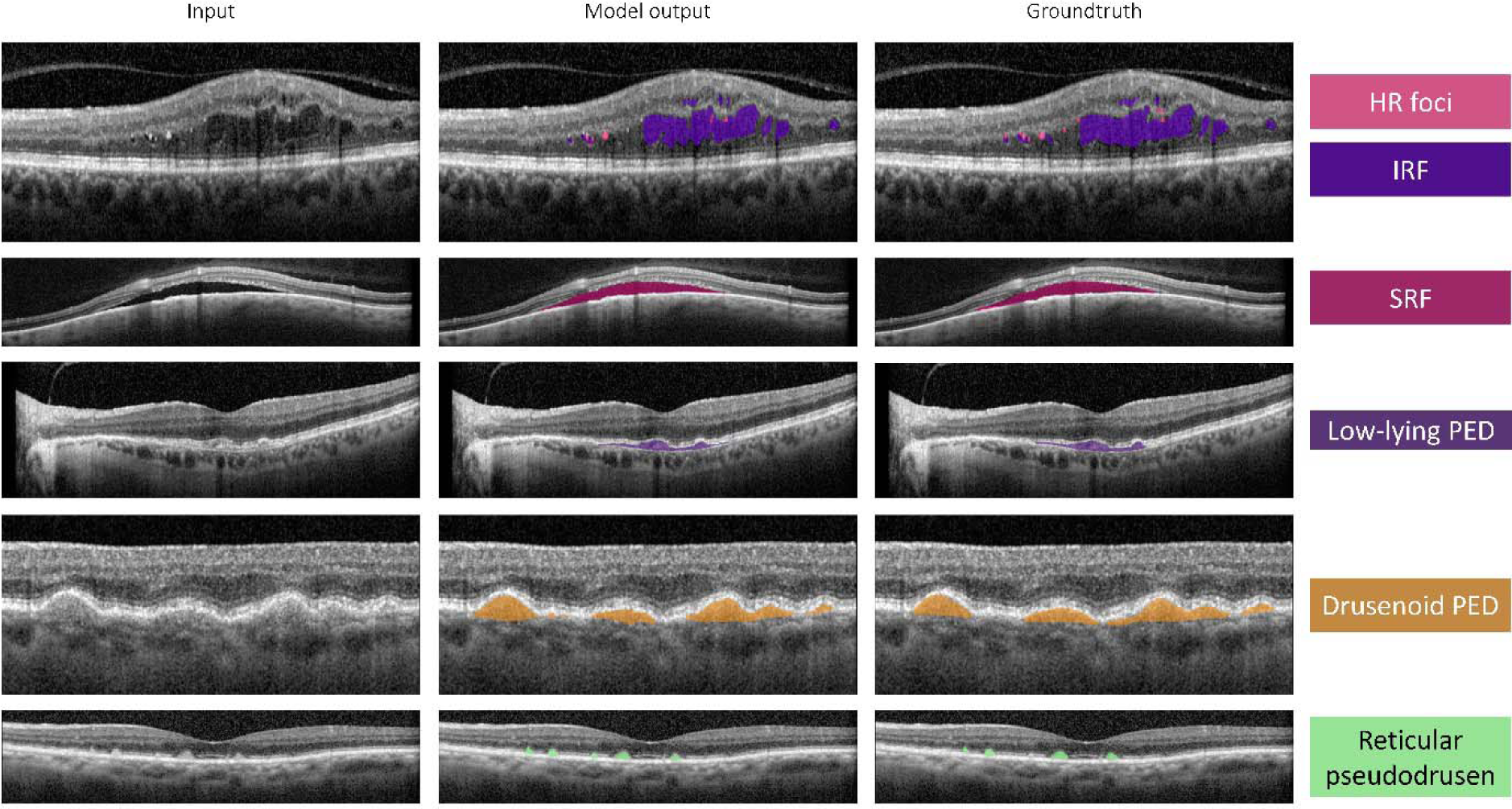
Comparison of MAYOCTransformer segmentation output with groundtruth segmentation masks of Hf foci, IRF, SRF, low-lying PED, drusenoid PED, and reticular pseudodrusen. Input OCT images (first column) with model segmentation (second column), and groundtruth segmentation (third column).

### Model testing against manual labelling variability

Total pixel area difference between model segmentation and ground truth segmentation was compared with total pixel area difference due to manual label-relabel variability for a separate hold-out test set. There was no statistical difference between mean absolute area error of automated segmentation and manual labeling variability for any of the discrete findings, except for choroid vessel segmentation, where model mean absolute area difference tended to be higher than manual labelling variability (**Table S2**).

### Model validation using external datasets and comparison against open-source models

When testing retinal fluid segmentation on an external annotated dataset [66], our segmentation model achieved Dice coefficient of 0.80. However, as shown in **Figure S5**, the original IRF segmentation maps contained many instances of missed IRF regions by the original annotators, which might have falsely decreased the Dice coefficient. When testing OCT layer segmentation on an externally annotated dataset (OCT5K) [67] without finetuning, our model achieved Dice coefficients that were consistently higher than the open-weight ReLayNet CNN-based neural network when tested head-to-head on the same external annotated dataset (**Table S3**).

Generalizability of choroid stroma and vessel segmentation was tested using publicly available swept-source OCT (ss-OCT) images. **Figure 8** presents randomly chosen images from the testing ss-OCT images. As shown in **Figure 8**, the open-weight CNN-based Choroidalyser model by Engelmann et al. [8] failed to produce accurate segmentation. The model either missed large portions of the choroid (**Figure 8A**, **C**, **D**, and **F**), hallucinated non-existent choroid and vessels (**Figure 8E**), or produced empty segmentation maps (**Figure 8B**). However, the model did produce very accurate choroid vessel and stroma segmentation with images from the training dataset presented by the authors (**Figure S6**). In contrast, MAYOCTransformer showed none of these issues and much better generalizability, as well as significantly higher Dice coefficients (0.69 [stroma] and 0.71 [vessels]) compared to Choroidalyser (0.46 [stroma] and 0.42 [vessels]). **Figure S7** further demonstrates the superiority of our model in generalizability using randomly selected images from the dataset by Kermany et al. [68].

**Figure 8:**
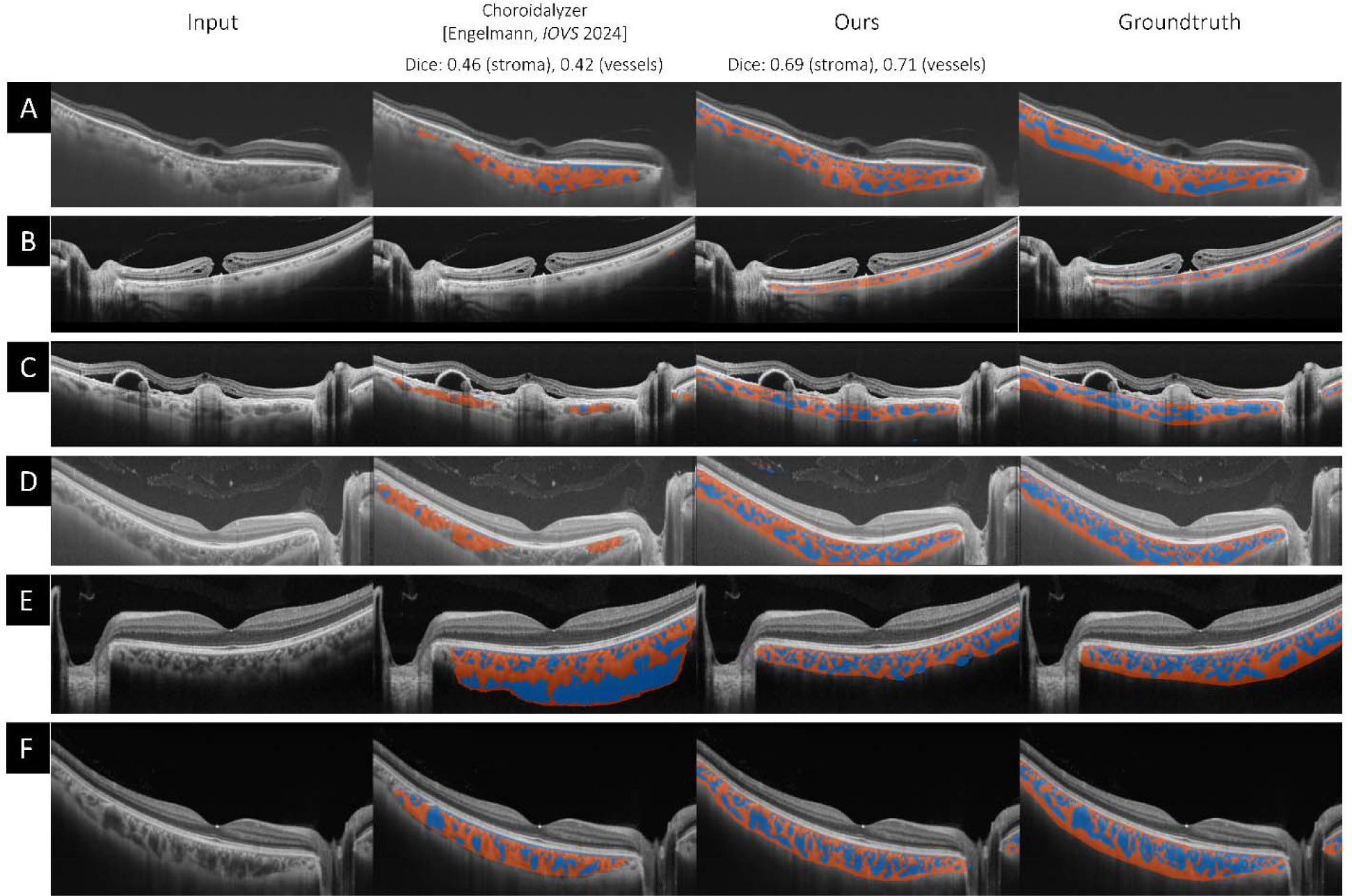
Comparison of MAYOCTransformer and Choroidalyser on swept-source OCT (ss-OCT) images. Representative test images from publicly available ss-OCT images (A–F). Original OCT images (First column). Segmentation predictions from Choroidalyser model showing various failure modes: undersegmentation, hallucinated structures, and empty masks (Second column). MAYOCTransformer predictions, which consistently segment both choroid stroma and vessel regions accurately across all examples (Third column). Ground truth segmentation masks (Forth column).

### Randomized blinded testing for segmentation model validation

As shown in **Table S4**, model segmentations obtained scores that were noninferior to those from manual segmentations. Interestingly, model scores were significantly higher for IRF, PED, Pseudodrusen, HR foci, ERM, and RNFL, while no statistically significant differences were found for all other segmentation classes. In addition, inter-rater agreement tended to be high (over 0.9) for answers for all segmentation classes except SHRM and Drusen (0.5 and 0.7, respectively).

### Model deployment

When performing inference on a single NVIDIA RTX 6000 Ada 48GB, the MAYOCTransformer models reached an average speed of 45.0 images per second. We deployed our models in our department’s retina service to generate automated reports on IRF, SRF, and PED volumes and geographic atrophy progression using a 97 b-scan protocol; models deployed on a single NVIDIA A2 GPU reached an average speed of 4.3 images per second, which allowed automated reports to be generated in under 3 minutes.

## DISCUSSION

This study presents MAYOCTransformer, the first transformer-based deep learning model for comprehensive semantic segmentation of retinal optical coherence tomography (OCT) images. We systematically compared its performance against several established convolutional neural network (CNN) architectures, including U-Net, U-Net++, FPN, and DeepLabV3+, across multiple segmentation tasks, including retinal layer segmentation, choroid segmentation, and the identification of discrete pathological findings. Our results demonstrate that MAYOCTransformer consistently outperformed CNN-based models across most segmentation tasks, particularly for retinal layers such as RNFL, GCL, and ERM, and for pathological findings such as fibrovascular pigment epithelial detachment (fv PED), subretinal hyper-reflective material (SHRM), and reticular pseudodrusen.

The MAYOCTransformer model achieved high Dice similarity coefficients (DSC) across all retinal layers, with significant improvements over CNN models for RNFL, GCL, and ERM segmentation. This suggests that the transformer-based architecture effectively captures global dependencies within OCT images, which may be particularly beneficial for structures that exhibit considerable morphological variation, such as the ERM. However, for certain layers, such as the photoreceptor outer segment (PR-OS) and retinal pigment epithelium (RPE), the U-Net model performed slightly better, indicating that for some anatomical structures, local spatial dependencies captured by CNNs remain competitive.

For choroid segmentation, MAYOCTransformer showed marginal improvements over CNN-based models, achieving slightly higher DSC values for both choroid stroma and choroid vessels. However, the differences in performance were not statistically significant, indicating that while transformers offer advantages in generalization, CNN models may still provide comparable segmentation accuracy for certain well-defined anatomical boundaries.

MAYOCTransformer demonstrated notable improvements in segmenting discrete pathological findings, particularly for fv PED, SHRM, and reticular pseudodrusen, where performance gains over CNN models were statistically significant. These findings suggest that transformer models may be better suited for capturing the complex, irregular morphologies of pathological features in OCT images. However, for certain fluid-related abnormalities such as intraretinal fluid (IRF) and subretinal fluid (SRF), the performance gains were more modest, and the differences in DSC values between MAYOCTransformer and CNN-based models did not reach statistical significance.

When applied to publicly available datasets, MAYOCTransformer demonstrated higher DSC values compared to open-weight CNN-based models, including ReLayNet for retinal layer segmentation and Choroidalyser for choroid segmentation. Notably, the Choroidalyser model failed to generalize to unseen data, producing erroneous or empty segmentation masks for several test images, whereas MAYOCTransformer maintained consistent segmentation accuracy across different imaging conditions.

Despite the promising results, this study has several limitations. First, our dataset was derived from a single institution (Mayo Clinic), which may limit the generalizability of our findings to OCT images acquired from different manufacturers or imaging protocols. Although external validation datasets were used, a broader evaluation using multi-center datasets would be beneficial.

## CONCLUSION

This study demonstrates that transformer-based architectures, specifically MAYOCTransformer, offer significant improvements over CNN-based models for the semantic segmentation of OCT images, particularly for retinal layers and discrete pathological findings. The model exhibits strong generalizability to external datasets and was non-inferior by expert graders in a blinded comparison study. These findings support the potential integration of transformer-based segmentation models into clinical ophthalmology workflows, paving the way for automated, high-precision analysis of OCT images.

