## Supplemental materials for "MAYOCTransformer: Masked-Attention for Yielding Comprehensive Semantic Segmentation of Retinal Optical Coherence Tomography Images using Transformer-based Neural Networks"

**Supplementary Table 1**: Basic image data characteristics of the complete OCT image database and the training subset

| | **Complete OCT database** | **Training image subset** | ***p-value*** |
|---|---|---|---|
| Total images | 18,234,749 | 3,500 | |
| Volumetric : radial images | 16,200,711 : 2,034,038 | 3,112 : 388 | 0.9 |
| Image height | 495.4 ± 0.003 | 495.4 ± 0.2 | 1.0 |
| Image width | 705.7 ± 0.08 | 708.4 ±5.6 | 0.6 |
| Mean pixel intensity | 44.1 ± 0.003 | 44.1 ± 0.18 | 0.9 |
| Signal-to-noise ratio | 0.82 ± 0.00004 | 0.82 ± 0.003 | 0.7 |

Values are presented as mean ± SEM. P-values are for independent samples t-tests.

**Supplementary Table 2**: Comparison of mean absolute area error of automated segmentation with manual labeling variability

| | **Model error (pixels)** | **Labeling variability (pixels)** | **p-value** |
|---|---|---|---|
| | | **Layers** | |
| **RNFL** | 567 ± 139 | 713 ± 196 | 0.9 |
| **GCL** | 476 ± 130 | 567 ± 139 | 0.9 |
| **IPL** | 351 ± 99 | 326 ± 85 | 0.4 |
| **INL** | 414 ± 69 | 393 ± 67 | 0.4 |
| **OPL** | 359 ± 65 | 387 ± 69 | 0.2 |
| **ONL** | 534 ± 98 | 524 ± 93 | 0.6 |
| **PR IS** | 282 ± 66 | 256 ± 64 | 0.3 |
| **PR OS** | 626 ± 74 | 593 ± 71 | 0.1 |
| **RPE** | 372 ± 50 | 344 ± 55 | 0.2 |
| **ERM** | 91 ± 34 | 106 ± 43 | 0.5 |
| | | **Choroid** | |
| **Stroma** | 2662 ± 621 | 2798 ± 612 | 0.4 |
| **Vessel** | 2635 ± 742 | 2374 ± 722 | 0.02 |
| | | **Discrete findings** | |
| **IRF** | 46.2 ± 19.5 | 42.6 ± 14.7 | 0.7 |
| **SRF** | 96.3 ± 60.9 | 76.8 ± 59.2 | 0.2 |
| **Drusenoid PED** | 7.6 ± 2.6 | 6.8 ± 2.8 | 0.2 |
| **Fv PED** | 260.0 ± 156.7 | 269.5 ± 228.9 | 0.9 |
| **Serous PED** | 8.2 ± 4.9 | 0.4 ± 0.2 | 0.1 |
| **SHRM** | 34.2 ± 11.6 | 25.0 ± 9.2 | 0.1 |
| **HR foci** | 15.1 ± 4.3 | 12.1 ± 3.1 | 0.2 |
| **Low-lying PED** | 56.8 ± 17.1 | 74.1 ± 21.3 | 0.1 |
| **Reticular pseudodrusen** | 25.3 ± 8.2 | 22.0 ± 7.4 | 0.3 |

Values are presented as mean ± SEM. p-values are for paired samples t-test.

**Supplementary Table 3**: External validation of OCT layer segmentation model and head-to-head comparison against ReLayNet using the OCT5K dataset

| | OCTransformer | ReLayNet |
|---|---|---|
| **Layer** | **Dice coefficient** | **Dice coefficient** |
| ILM to INL | **0.87** | 0.82 |
| OPL to PR-OS | **0.82** | 0.78 |
| PR-OS | **0.82** | 0.78 |
| RPE | **0.74** | 0.65 |

ILM: internal limiting membrane; INL: inner nuclear layer; OPL: outer plexiform layer; PR-OS: photoreceptor-outer segment; RPE: retinal pigment epithelium.

**Supplementary Table 4:** Randomized blinded testing for segmentation model validation

| Class | Model score (0-4) | Manual score (0-4) | p-value | Inter-rater agreement |
|---|---|---|---|---|
| **IRF** | 2.9 | 1.1 | 0.01 | 0.9 |
| **SRF** | 2.3 | 1.7 | 0.3 | 0.9 |
| **PED** | 3.3 | 0.8 | 0.04 | 1.0 |
| **Drusen** | 2.1 | 1.8 | 0.7 | 0.7 |
| **Pseudodrusen** | 3.5 | 0.2 | 0.001 | 0.9 |
| **SHRM** | 2.0 | 2.0 | 1.0 | 0.5 |
| **HR foci** | 3.3 | 0.8 | 0.05 | 1.0 |
| **ERM** | 2.8 | 1.2 | 0.05 | 0.9 |
| **RNFL** | 2.4 | 1.6 | 0.05 | 1.0 |
| **GCL** | 2.3 | 1.7 | 0.07 | 1.0 |
| **IPL** | 2.0 | 1.9 | 0.7 | 1.0 |
| **INL** | 2.1 | 1.9 | 0.7 | 1.0 |
| **OPL** | 2.1 | 1.9 | 0.7 | 0.9 |
| **ONL** | 2.2 | 1.8 | 0.4 | 0.9 |
| **PR-IS** | 2.4 | 1.6 | 0.09 | 1.0 |
| **PR-OS** | 2.4 | 1.6 | 0.1 | 0.9 |
| **RPE** | 2.3 | 1.7 | 0.1 | 0.9 |
| **Choroid stroma** | 2.0 | 2.0 | 1.0 | 1.0 |
| **Choroid vessel** | 2.0 | 2.0 | 1.0 | 1.0 |

Scores are reported as mean (values range from 0 to 4). P-values are for paired samples t-test.

**Figure S1**: Online quiz-based randomized blinded testing

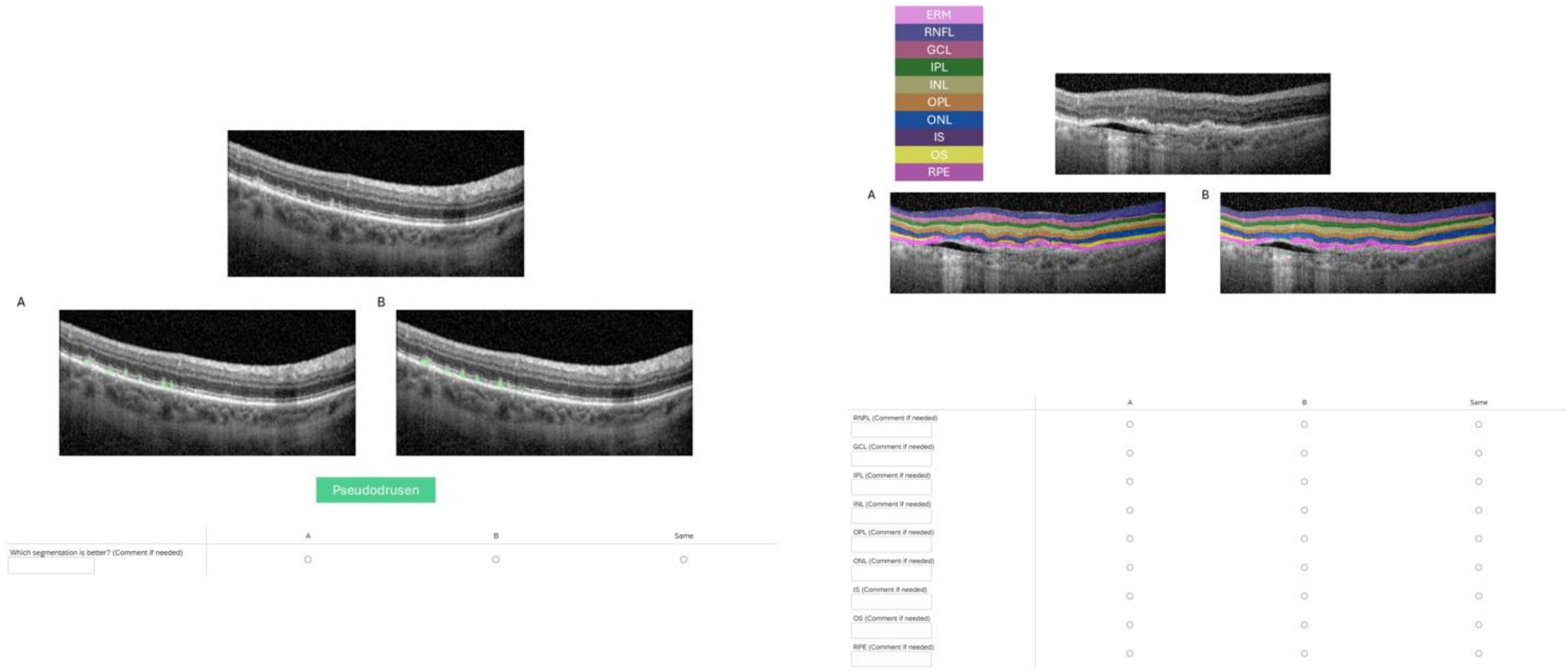

**Figure S2**: Comparison of dice coefficients of segmentation models for retinal layer segmentation

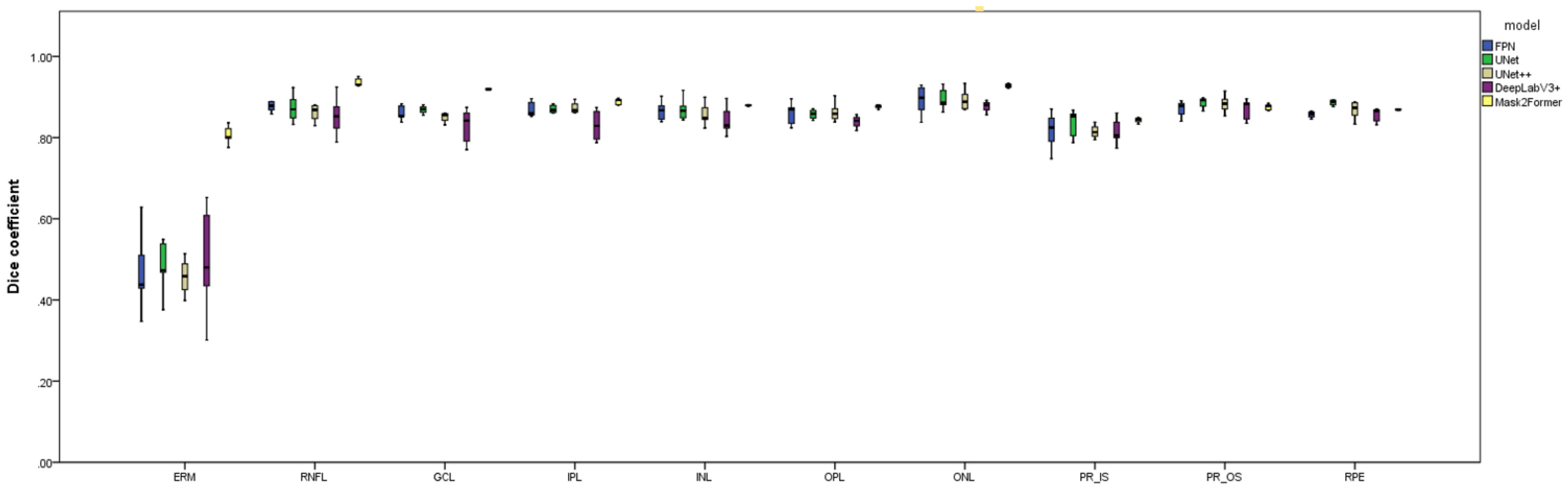

**Figure S3**: Comparison of dice coefficients of segmentation models for choroid segmentation

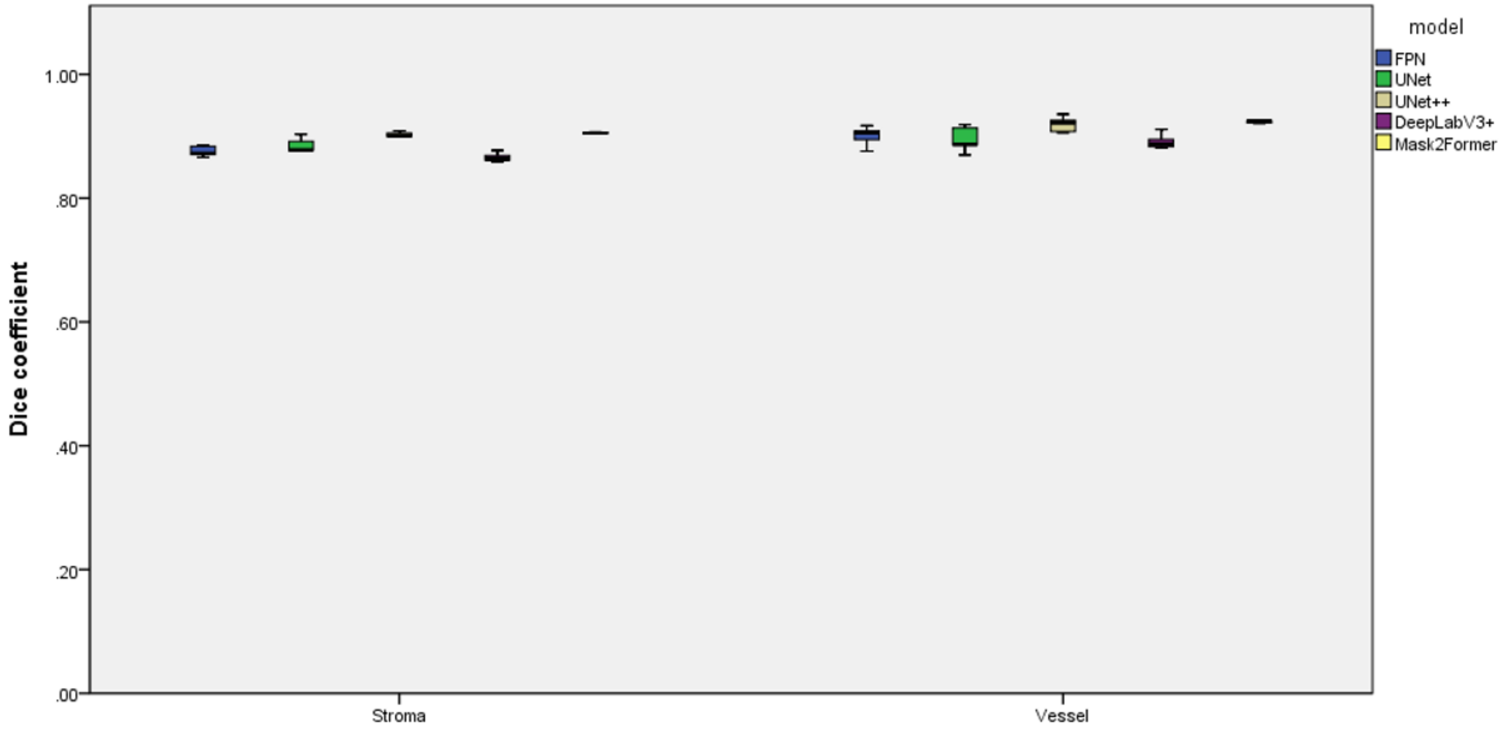

**Figure S4**: Comparison of dice coefficients of segmentation models for discrete finding segmentation

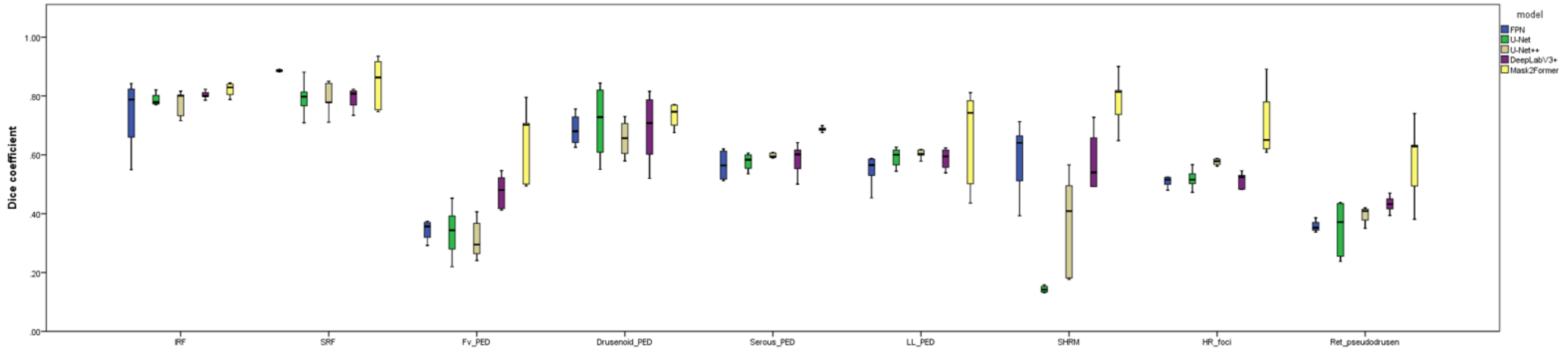

**Figure S5**: Comparison of intraretinal fluid segmentation with ground truth segmentation in an external dataset

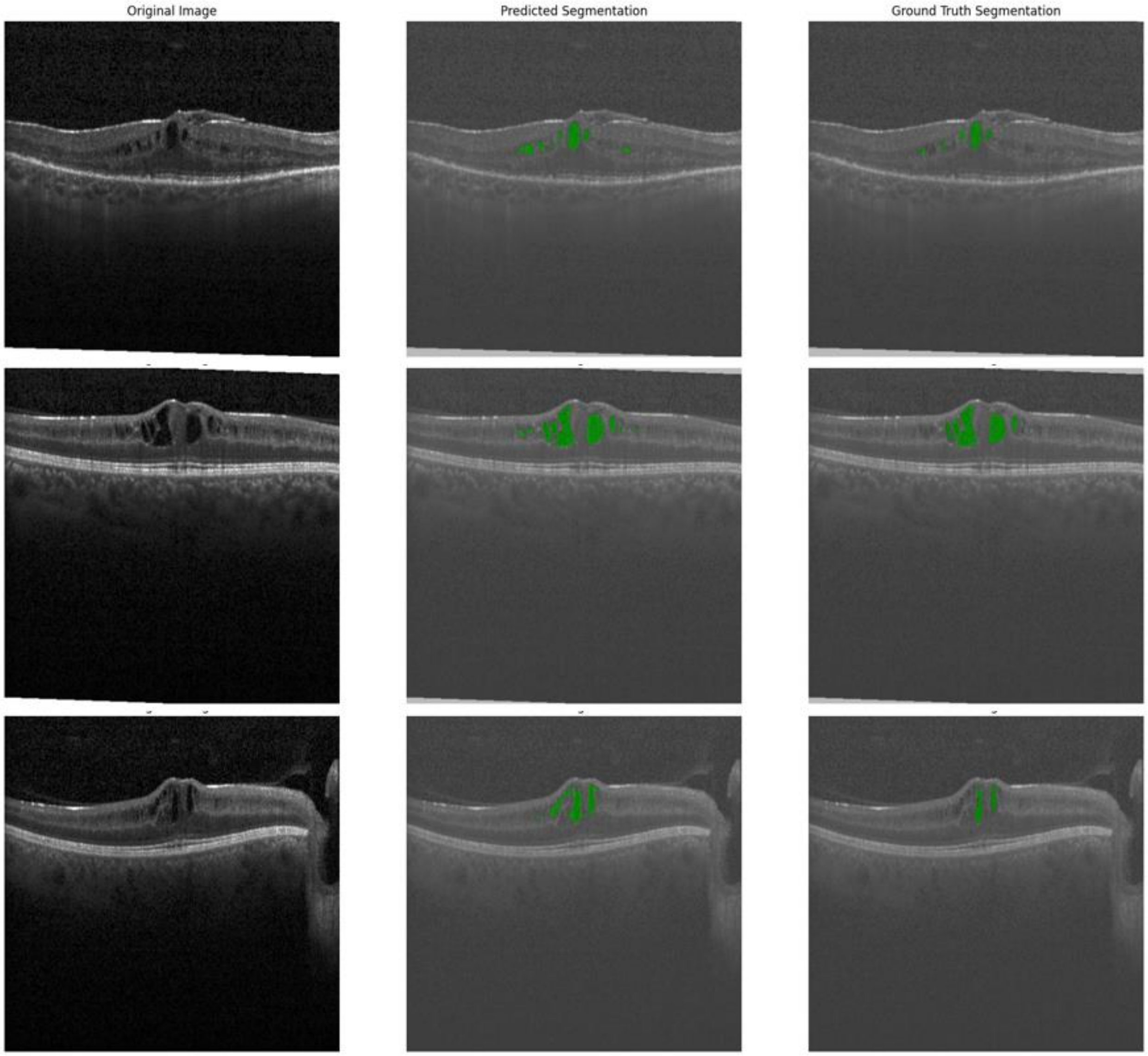

**Figure S6**: Model outputs of Choroidalyzer using in-distribution input image

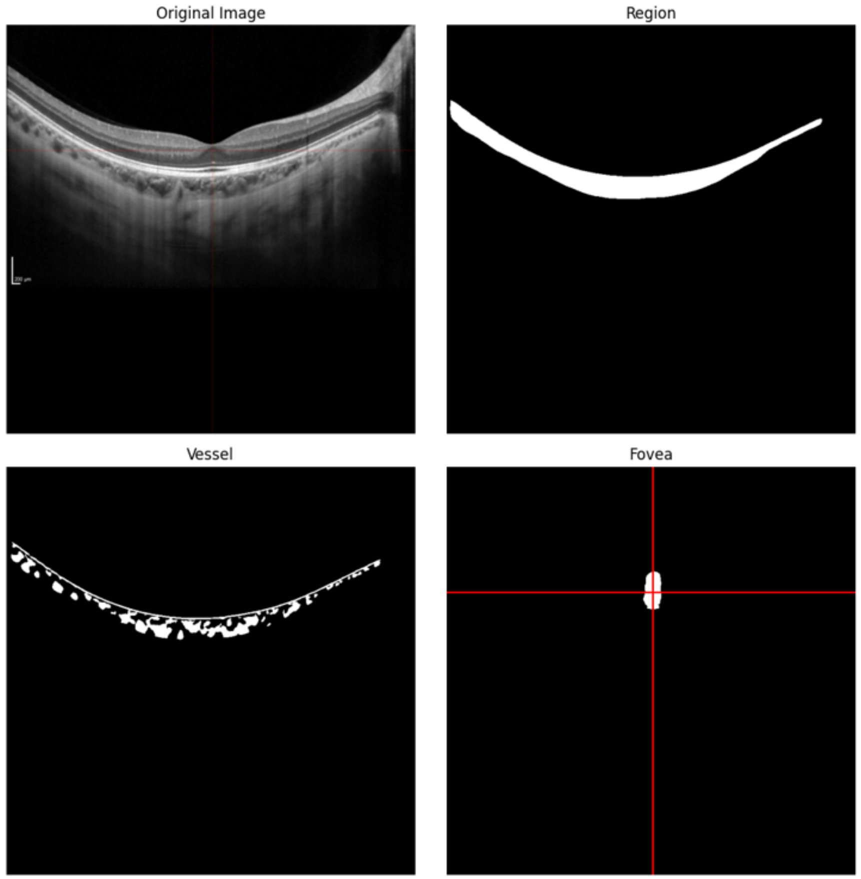

**Figure S7**: Comparison of model outputs between OCTransformer and Choroidalyzer using out-of-distribution input images

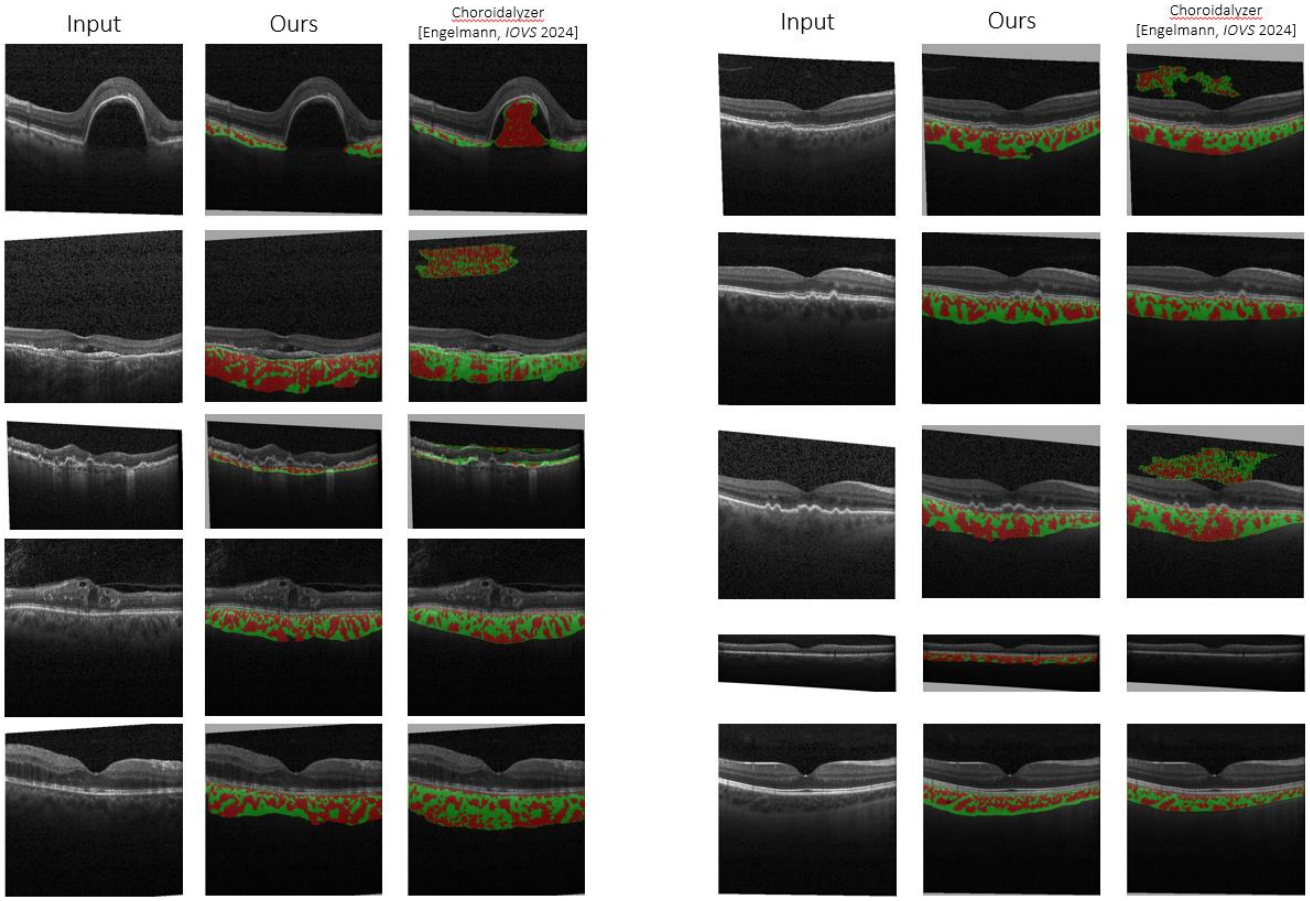
